# The drift diffusion model as the choice rule in inter-temporal and risky choice: a case study in medial orbitofrontal cortex lesion patients and controls

**DOI:** 10.1101/642587

**Authors:** Jan Peters, Mark D’Esposito

## Abstract

Sequential sampling models such as the drift diffusion model have a long tradition in research on perceptual decision-making, but mounting evidence suggests that these models can account for response time distributions that arise during reinforcement learning and value-based decision-making. Building on this previous work, we implemented the drift diffusion model as the choice rule in inter-temporal choice (*temporal discounting*) and risky choice (*probability discounting*) using a hierarchical Bayesian estimation scheme. We validated our approach in data from nine patients with focal lesions to the ventromedial prefrontal cortex / medial orbitofrontal cortex (vmPFC/mOFC) and nineteen age- and education-matched controls. Choice model parameters estimated via standard softmax action selection were reliably reproduced using the drift diffusion model as the choice rule, both for temporal discounting and risky choice. Model comparison revealed that, for both tasks, the data were best accounted for by a variant of the drift diffusion model including a non-linear mapping from value-differences to trial-wise drift rates. Posterior predictive checks of the winning models revealed a reasonably good fit to individual participants reaction time distributions. We then applied this modeling framework and 1) reproduced our previous results regarding temporal discounting in vmPFC/mOFC patients and 2) showed in a previously unpublished data set on risky choice that vmPFC/mOFC patients exhibit increased risk-taking relative to controls. Analyses of diffusion model parameters revealed that vmPFC/mOFC damage abolished neither value sensitivity nor asymptote of the drift rate. Rather, it substantially increased non-decision times and reduced response caution during risky choice. Our results highlight that novel insights can be gained from applying sequential sampling models in studies of inter-temporal and risky decision-making in cognitive neuroscience.

## Introduction

Understanding the neuro-cognitive mechanisms underlying decision-making and reinforcement learning^1–3^ has potential implications for many neurological and psychiatric disorders associated with maladaptive choice behavior^4–6^. Modeling work in value-based decision-making and reinforcement learning often relies on simple logistic (softmax) functions^7,8^ to link model-based decision values to observed (often binary) choices. In contrast, in perceptual decision-making, sequential sampling models such as the drift diffusion model (DDM) that not only account for the observed choices but also for the full reaction time (RT) distributions have a long tradition^9–11^. Recent work in reinforcement learning^12–15^, inter-temporal choice^16,17^ and simple value-based choice^18–21^ has shown that sequential sampling models can be successfully applied in these domains.

In the DDM, decisions arise from a noisy evidence accumulation process that terminates as the accumulated evidence reaches one of usually two response boundaries^9^. In cases when there is an objectively correct response such as during perceptual decision-making, the upper boundary typically codes correct responses and the lower boundary codes errors, a coding scheme henceforth referred to as *accuracy coding*. In its simplest form, the DDM has four free parameters: the boundary separation parameter *α* governs how much evidence is required before committing to a decision. In accuracy coding, the boundary separation parameter therefore reflects the speed-accuracy trade-off: a lower boundary separation leads to faster but more error-prone responses, whereas a greater boundary separation leads to slower but more accurate responses. The drift rate parameter *v* determines the mean rate of evidence accumulation during a trial. For accuracy coding, a greater drift rate reflects a greater rate of evidence accumulation and thus faster and more accurate responding. In contrast, a drift rate of zero would indicate chance level performance, as the evidence accumulation process would have an equal likelihood of terminating at the correct or error boundaries (for a neutral bias). Finally, the starting point or bias parameter *z* determines the starting point of the evidence accumulation process in units of the boundary separation, and the non-decision time *τ* reflects components of the reaction time related to stimulus encoding and/or response preparation that are unrelated to the evidence accumulation process. The DDM can account for a wide range of experimental effects on RT distributions during simple choices^9^.

Recent studies on reinforcement learning have similarly used accuracy coding when fitting the DDM^14,15^ to describe how choices and RT distributions relate to learned action values. In these studies, the upper boundary was defined as a selection of the stimulus with the objectively better reinforcement rate, and the lower boundary as a selection of the objectively inferior stimulus. However, while in perceptual decision tasks all information required to make a correct response is in principle available to the participant on any given trial, this is not necessarily the case in reinforcement learning. Here, a correct evaluation of the optimal stimulus depends also on the experienced reinforcement history. For situations without an objectively correct response, DDM boundaries have sometimes been coded in a way that reflects the degree to which choices are consistent with e.g. preference ratings^18^ or monetary bids^22^ collected before the decision phase. Accuracy in such a scenario then corresponds to choice consistency. However, this approach is not feasible when the goal is to use the DDM to estimate the very preferences that in this approach are used to implement accuracy coding. Alternatively, one could abandon the idea of “accuracy” altogether. This is the approach that was taken in the present study. Instead of accuracy coding, we used a *stimulus coding* scheme, such that each boundary in the DDM corresponds to a different choice category, e.g. “face” vs. “house” in a perceptual decision task, “choice of the risky option” vs. “choice of the safe option” in a risky choice setting. This also allows one to estimate a response bias towards one of the decision boundaries.

The application of sequential sampling models such as the DDM has several potential advantages over traditional softmax^7^ choice rules. First, including RT data during model estimation may improve both the reliability of the estimated parameters^12^ and parameter recovery^13^, thereby leading to more robust estimates. Second, taking into account the full RT distributions can reveal additional information regarding the dynamics of decision processes^14,15^. This is of potential interest, in particular in the context of maladaptive behaviors in clinical populations^14,23–26^ but also when the goal is to more fully account for how decisions arise on a neural level^10^.

In the present case study, we focus on a brain region that has long been implicated in decision-making, reward-based learning and impulse regulation^27,28^, the ventromedial prefrontal / medial orbitofrontal cortex (vmPFC/mOFC). Performance impairments on the Iowa Gambling Task are well replicated in vmPFC/mOFC patients^27,29,30^. Damage to vmPFC/mOFC also increases temporal discounting^31,32^ (but see^33^) and risk-taking^34–36^, impairs reward-based learning^37–39^ and has been linked to inconsistent choice behavior^40–42^. Meta-analyses of functional neuroimaging studies strongly implicate this region in reward valuation^43,44^. Based on these observations, we reasoned that vmPFC/mOFC damage might also render RTs during decision-making less dependent on value. In the context of the DDM, this could be reflected in changes in the value-dependency of the drift rate v. In contrast, more general impairments in the processing of decision options, response execution and/or preparation would be reflected in changes in the non-decision time. Interestingly, however, one previous model-free analysis in vmPFC/mOFC patients revealed a similar modulation of RTs by value in patients and controls^41^.

The present study therefore had the following aims. The first aim was a validation of the applicability of the DDM as a choice rule in the context of inter-temporal and risky choice. To this end, we first performed a model comparison of variants of the DDM in a data set of nine vmPFC/mOFC lesion patients and nineteen controls and performed a number of model validation tests. Second, since recent work on reinforcement learning suggested that the mapping from value differences to trial-wise drift rates might be non-linear^15^ rather than linear^14^, we compared these different variants of the DDM in our data and ran posterior predictive checks on the winning DDM models to explore the degree to which the observed RT distributions could be accounted for by the best-fitting models. Third, we re-analyzed previously published temporal discounting data in controls and vmPFC/mOFC lesion patients to examine the degree to which our previously reported model-free analyses^31^ could be reproduced using a hierarchical Bayesian model-based analysis with the DDM as the choice rule. Fourth, we used the same modeling framework to analyze previously unpublished data from a risky decision-making task in the same lesion patients and controls to examine whether risk taking in the absence of a learning requirement is increased following vmPFC/mOFC damage. Finally, we explored changes in choice dynamics as revealed by DDM parameters as a result of vmPFC/mOFC lesions.

## Methods

### Procedure

We report data from two value-based decision-making tasks: one previously unpublished data from a risky-choice task and one previously published data set from a temporal discounting task (see below for task details). Data were acquired in nine patients with focal lesions that included medial orbitofrontal cortex and nineteen healthy age- and education-matched controls. The temporal discounting task was always performed first, followed by the risky choice task.

For a detailed account of etiology and socio-demographic information for all participants, the reader is referred to our previous paper^31^. For convenience, we reproduce the lesion overlap plot from our previous paper in Figure 1. All participants gave informed written consent, and the study procedure was approved by the local institutional review board of the University of California, Berkeley, USA.

**Figure 1.**
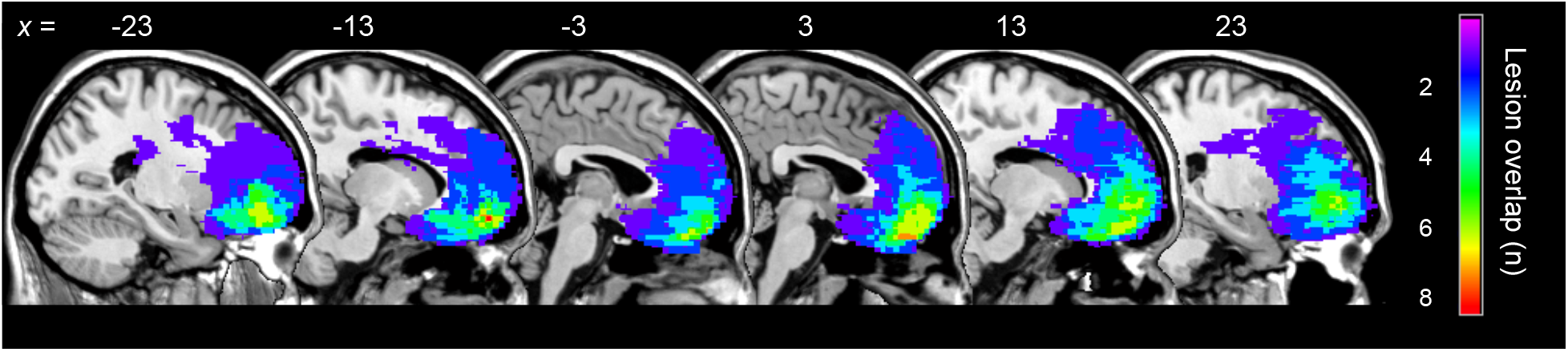
Lesion overlap in the vmPFC/mOFC lesion patient group reproduced from Peters & D’Esposito (2016)^31^. The color code reflects the number of patients with overlapping lesions in each voxel.

#### Temporal discounting task

Here participants performed 224 trials of an inter-temporal choice task involving a series of choices between smaller-but-sooner (SS) and larger-but-later (LL) rewards. On half the trials, the SS reward was available immediately (*now* condition), whereas on the other half of the trials, the SS reward was available only after a 30d delay (*not now* condition). In the *now* condition, the SS reward consisted of $10 available immediately and LL rewards consisted of all combinations of fourteen reward amounts (10.1, 10.2, 10.5, 11, 12, 15, 18, 20, 30, 40, 70, 100, 130, 150 dollars) and seven delays (1, 3, 5, 8, 14, 30, 60 days). Trials for the *not now* condition where identical, with the exception that an additional delay of 30 days was added to both options, such that in *not now* trials, the SS reward was always associated with a 30 day delay, and LL reward delays ranged from 31 to 91 days. Trials were presented in randomized order and with a randomized assignment of options to the left/right side of the screen. Options remained on the screen until a response was logged.

#### Risky choice task

Here participants made a total of 112 choices between a certain (100% probability) $10 reward and larger-but-riskier options. The risky options consisted of all combinations of sixteen reward amounts (10.1, 10.2, 10.5, 11, 12, 15, 18, 20, 25, 30, 40, 50, 70, 100, 130, 150 dollars) and seven probabilities (10%, 17%, 28%, 54%, 84%, 96%, 99%). Trials were presented in randomized order and with a randomized assignment of options to the left/right side of the screen. As in the temporal discounting task, options remained on the screen until a response was logged.

Participants were instructed that all choices from the two tasks were potentially behaviorally relevant. A single trial was pseudo-randomly selected following completion of both tasks, and participants received their choice from that trial as a cash bonus.

### Temporal discounting model

Based on previous work on the effect of SS immediacy on discounting behavior ^45,46^, we hypothesized discounting to be hyperbolic relative to the soonest available reward. Previous studies^31,46^ fitted separate discount rate parameters to trials with immediate vs. delayed SS rewards. Here we extended this approach by instead fitting a single k-parameter (reflecting discounting in the *now* condition), and a subject-specific shift parameter *s* modeling the reduction in log(k) in the *not now* condition as compared to the *now* condition:

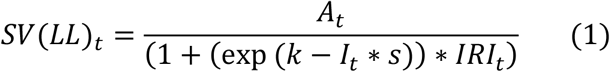

Here, *SV* is the subjective discounted value of the delayed rewards, *A* is the amount of the *LL* reward on trial t, *k* is the subject specific discount rate for *now* trials in log-space, *I* is an indicator variable coding the condition (0 for *now* trials, 1 for *not now* trials), *s* is a subject-specific shift in log(*k*) between *now* and *not-now* conditions and *IRI* is the inter-reward-interval on trial *t*. Note that this model corresponds to the elimination-by-aspects model of Green et al. ^45^.

### Risky choice model

Here we applied a simple one-parameter probability discounting model^47,48^, where discounting is hyperbolic over the odds-against-winning the gamble:

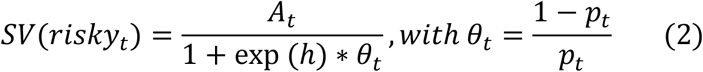

Here SV is the subjective discounted value of the risky reward, *A* is the reward amount on trial *t* and *θ* is the odds-against winning the gamble. The probability discount rate *h* (again fitted in log-space) models the degree of value discounting over probabilities. We also fit the data with a two-parameter model that includes separate parameters for the curvature and elevation of the probability weighting function ^49–51^. However, when fitting a 2-parameter model at the single subject level, in a number of individual cases the posterior distributions of the curvature and/or elevation parameters did not have a clear peak or a clearly Gaussian shape, suggesting that we likely did not have adequate coverage of the probability and amount dimensions to reliably dissociate these different components of risk preferences. For this reason we opted for the simpler single-parameter hyperbolic model instead.

### Softmax choice rule

Standard softmax action selection models the probability of choosing the LL reward (or the risky option) on trial *t* as:

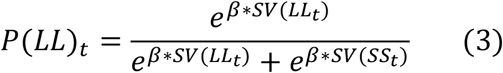

Here, *SV* is the subjective value of the LL reward according to Eq. 1 (or the risky reward according to Eq. 2) and *β* is an inverse temperature parameter, modeling choice stochasticity (for *β* = 0, choices are random and as *β* increases, choices become more dependent on the option values).

### Drift diffusion choice rule

For the DDMs, we build on earlier work in reinforcement learning^14,15^ and inter-temporal choice^13,16^. Specifically, we replaced the softmax action selection rule (see previous section) with the drift diffusion model as the choice rule, using the Wiener module^52^ for the JAGS software package^53^. In contrast to previous reinforcement learning approaches^14,15^ that used accuracy coding for the boundary definitions, we here used stimulus coding, such that the lower boundary (0) was defined as a selection of the SS reward (or the 100% option in the case of risky choice), and the upper boundary (1) as selection of the LL reward (or the risky option in the case of risky choice). This is because we were explicitly interested in modeling a bias towards SS vs. LL options. RTs for choices towards the lower boundary were multiplied by −1 prior to estimation.

We initially used absolute RT cut-offs for trial exclusion^14^ such that 0.4s < RT < 10s. However, when using such an absolute cut-off, single fast outlier trials can still force the non-decision-time to adjust to accommodate these observations, which can lead to a massive negative impact on model fit at the individual-subject level. This is also what we observed in two participants when plotting posterior predictive checks from hierarchical models with absolute cut-offs. For this reason, we instead excluded for each participant the slowest and fastest 2.5% of trials from analysis, which eliminated the problem. The reaction time on trial *t* is then distributed according to the Wiener first passage time (WFPT):

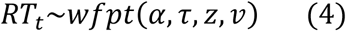

Here, *α* is the boundary separation (modeling response caution / the speed-accuracy trade-off), *z* is the starting point of the diffusion process (modeling a bias towards one of the decision boundaries), *τ* is the non-decision time (reflecting perceptual and/or response preparation processes unrelated to the evidence accumulation process) and *v* is the drift rate (reflecting the rate of evidence accumulation). Note that in the JAGS implementation of the Wiener model^52^, the starting point *z* is coded in relative terms and takes on values between 0 and 1. That is, *z* = .5 reflects no bias, *z* >.5 reflects a bias towards the upper boundary, and *z* <.5 a bias towards the lower boundary.

In a first step, we fit a null model (DDM_0_) that included no value modulation. That is, the null model for both the temporal discounting and risky choice data had four free parameters (*α*, τ, *v*, and *z*) that for each participant were constant across trials. Next, to link the diffusion process to the valuation models (Eq. 1, Eq., 2), we compared two previously proposed functions linking trial-by-trial variability in the drift rate *v* to value differences. First, we used a linear mapping as proposed by Pedersen et al. (2017)^14^:

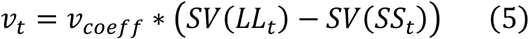

Here, *v*_*coeff*_ is a free parameter that maps value differences onto the drift rate *v* and simultaneously transforms value differences to the appropriate scale of the DDM^14^. This implementation naturally gives rise to the effect that highest conflict (when values are highly similar) would be expected to be associated with a drift rate close to zero. For positive values of *v*_*coeff*_, as *SV(SS)* increases over *SV(LL)*, the drift rate becomes more negative, reflecting evidence accumulation towards the negative (*SS*) boundary. The reverse is the case as *SV(LL)* increases over *SV(SS)*. For the risky choice models, *SV(LL)* was replaced with *SV(risky)*, and *SV(SS)* with *SV(safe)*. Second, we also applied an additional non-linear transformation of the scaled value differences via the S-shaped function suggested by Fontanesi et al. (2019)^15^:

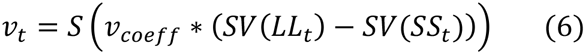

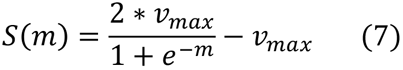

*S* is a sigmoid function centered at 0 with slope *m* and asymptote ± *v*_*max*_. Again, effects of choice difficulty on the drift rate naturally arise: for highest decision conflict when *SV*(*SS*) = *SV*(*LL*), the drift rate would again be zero, whereas for larger value differences, *v* increases up to a maximum of ±*v*_*max*_. Table 1 provides an overview of the parameters of the DDM_S_ model.

**Table 1.**
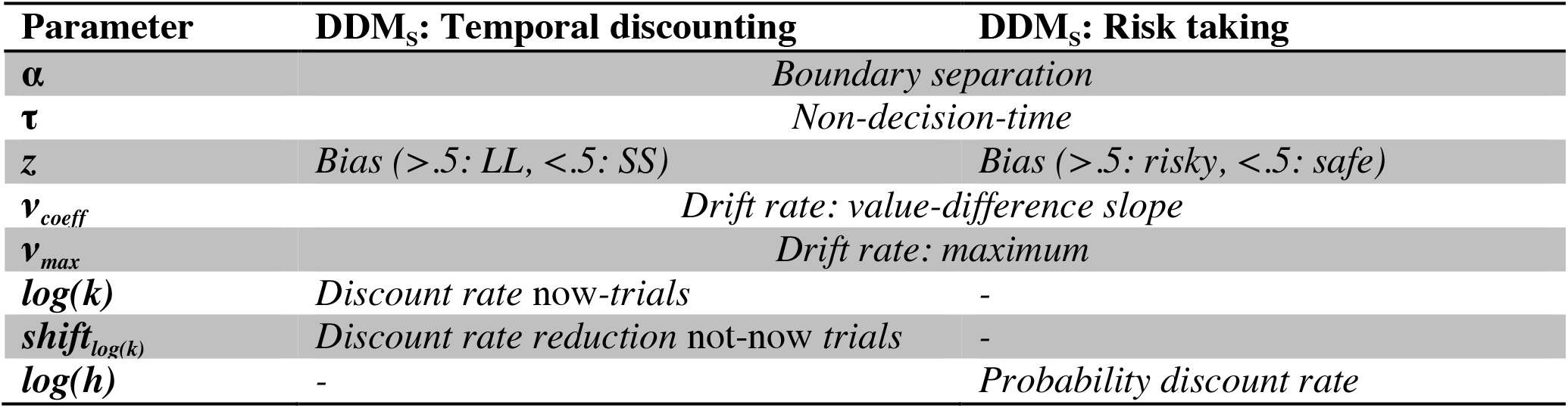
Overview of the parameters of the DDM_S_ models

### Hierarchical Bayesian models

We used the following model-building procedure. In a first step, models were fit at the single-subject level. After validating that reasonably good fits could be obtained for single-subject data (by ensuring that 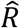 statistic was in an acceptable range of 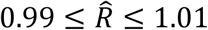 and the posterior distributions were clearly peaked and centered at reasonable parameter values) we re-fit all models using a hierarchical framework with separate group-level distributions for controls and mOFC patients. We again assessed chain convergence such that values of 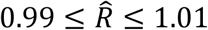 were considered acceptable for all group- and individual-level parameters. For the group-level hyperparameters, we used uninformative priors (e.g. uniform distributions for means defined over sensible ranges, gamma distributions for precision). All model code is available on the Open Science Framework (https://osf.io/5rwcu/).

### Model estimation and comparison

All models were fit using Markov Chain Monte Carlo (MCMC) as implemented in JAGS^53^ with the *matjags* interface (https://github.com/msteyvers/matjags) for Matlab © (The Mathworks) and the JAGS Wiener package^52^. For each model, we ran two chains with a burn-in period of 50k samples and thinning of 2. 10k further samples were then retained for analysis. Chain convergence was assessed via the 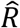 statistic, where we considered 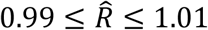 as acceptable values. Relative model comparison was performed via the Deviance Information Criterion (DIC), where lower values indicate a better fit^54^.

### Posterior predictive checks

Because a superior relative model fit does not necessarily mean that the winning model captures key aspects of the data, we additionally performed posterior predictive checks. To this end, during model estimation, we simulated 10k full datasets from the hierarchical models, based on the posterior distribution of the parameters. Model predicted RTs for a random sample of 1k of these simulated data sets were then smoothed via non-parametric density estimation in Matlab (*ksdensity.m*) and overlaid on the observed RT distributions for each individual participant.

### Analysis of group differences

To characterize group differences, we show posterior distributions for all parameters, as well as 85% and 95% highest density intervals for the difference distributions of the group posteriors. We furthermore report Bayes Factors for directional effects^14,55^ based on these difference distributions as *BF* = *i*/(1 − *i*) were *i* is the integral of the posterior distribution from 0 to +∞, which we estimated via non-parametric kernel density estimation in Matlab (*ksdensity.m*). Following common criteria^56^, Bayes Factors > 3 are considered positive evidence, and Bayes Factors > 12 are considered strong evidence. Bayes Factors < 0.33 are likewise interpreted as evidence in favor of the alternative model. Finally, we report standardized measures of effect size (Cohen’s *d*) which we calculated based on the mean posterior distributions of the group means and the pooled standard deviations across groups, which we calculated using the means of the group posterior distributions for the precision

### Data availability

Trial-wise behavioral data of all participants are available from the first author upon request.

### Code availability

JAGS model code for all models is available on the Open Science Framework (https://osf.io/5rwcu/).

## Results

### Model comparison

We first compared the fit of two previously proposed DDM models with linear (DDM_lin_, see Eq. 5)^14^ and non-linear (DDM_s_, see Eq. 6 and Eq. 7)^15^ value-dependent drift-rate scaling in terms of the deviance information criterion (DIC)^54^. For comparison we also included a null model (DDM_0_) with constant drift rate, that is, a model without value modulation. For both temporal discounting data (Table 2) and risky choice / probability discounting data (Table 3), the non-linear drift rate scaling models outperformed linear scaling, and both models fit the data better than the DDM_0_.

**Table 2.** Model comparison of drift diffusion models of temporal discounting. The hyperbolic+shift value function (see Eq. 1) corresponds to hyperbolic discounting in the *now* condition, and a shift parameter that models the decrease in discounting between the *now* and *not now* conditions.

| Model | Value scaling | Value function | DIC | Rank |
| --- | --- | --- | --- | --- |
| DDM<sub>0</sub> | - | - | 20889 | 3 |
| DDM<sub>lin</sub> | *Linear* | *Hyperbolic+Shift* | 19523 | 2 |
| DDM<sub>s</sub> | *Sigmoid* | *Hyperbolic+Shift* | 16845 | 1 |

**Table 3.** Model comparison of drift diffusion models of risky choice. The hyperbolic value function (see Eq. 2) corresponds to hyperbolic discounting over the odds-against-winning the gamble.

| Model | Value scaling | Value function | DIC | Rank |
| --- | --- | --- | --- | --- |
| DDM<sub>0</sub> | - | - | 11477 | 3 |
| DDM<sub>lin</sub> | *Linear* | *Hyperbolic* | 10281 | 2 |
| DDM<sub>s</sub> | *Sigmoid* | *Hyperbolic* | 9204.2 | 1 |

### Posterior predictive checks

While a relative model comparison is helpful to select among a set of candidate models, it is nonetheless important to verify that the winning model captures the overall pattern in the data, which we assessed via posterior predictive checks (see methods section). We plot posterior predictive checks for the reaction time distribution of each individual participant. Figures 2 and 3 show smoothed individual-participant RT distributions simulated from the posterior of the winning hierarchical models (DDM_S_) overlaid on the observed single-subject RT distributions for temporal discounting (Figure 2) and risky choice / probability discounting (Figure 3). As can be seen, the DDM_S_ provided a reasonable account of the observed RT distributions. However, RTs for SS choices were slightly underestimated in both groups

**Figure 2.**
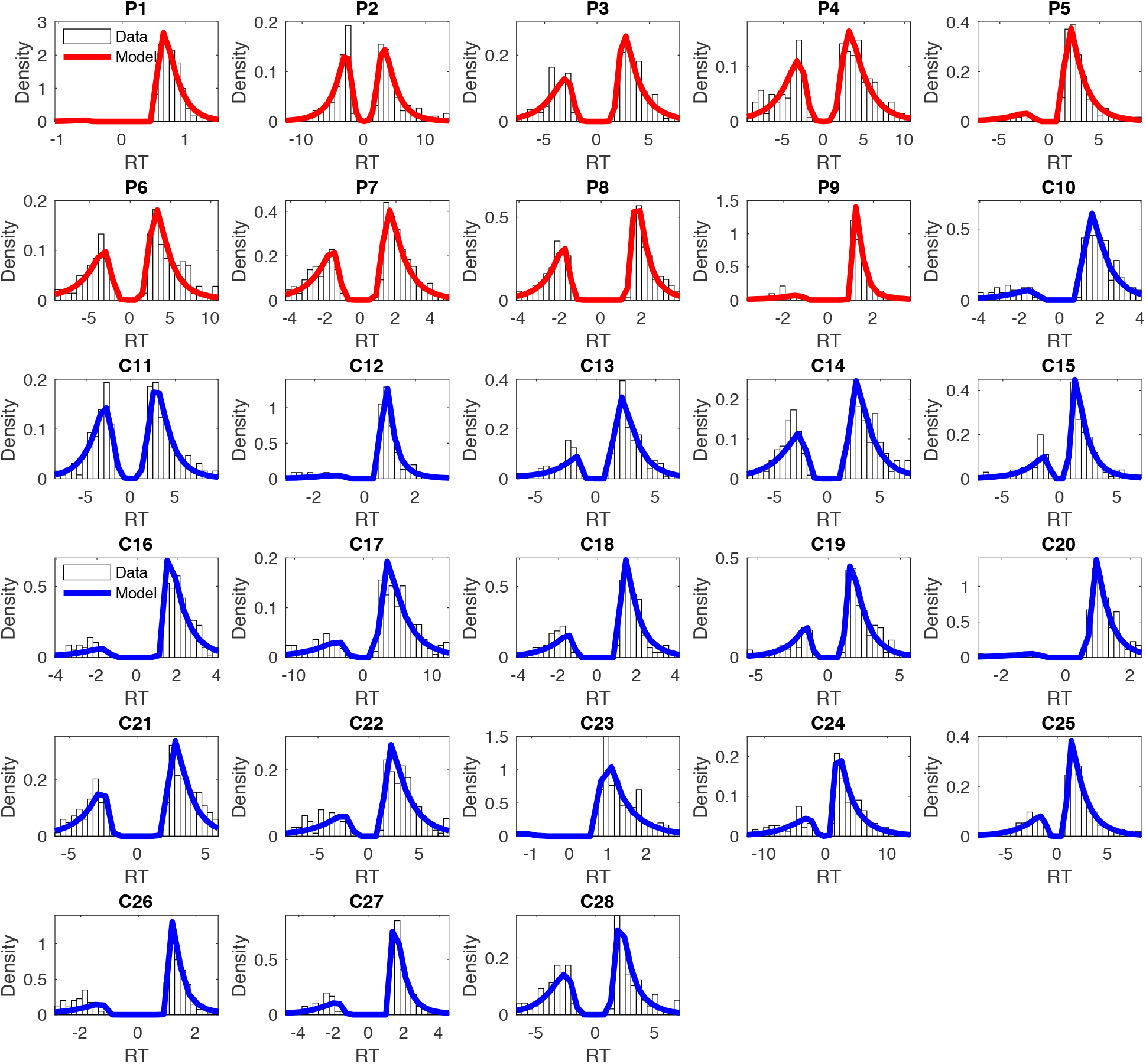
Posterior predictive plots of the drift diffusion temporal discounting model with non-linear value scaling of the drift rate (DDM_S_) for all participants (red – mOFC patients, blue – controls). Histograms depict the observed RT distributions for each participant. The solid lines are smoothed histograms of the model predicted RT distributions from 1000 individual subject data sets simulated from the posterior of the winning hierarchical model. RTs for smaller-sooner choices are plotted as negative, whereas RTs for larger-later choices are plotted as positive. The x-axes are adjusted to cover the range of observed RTs for each participant.

**Figure 3.**
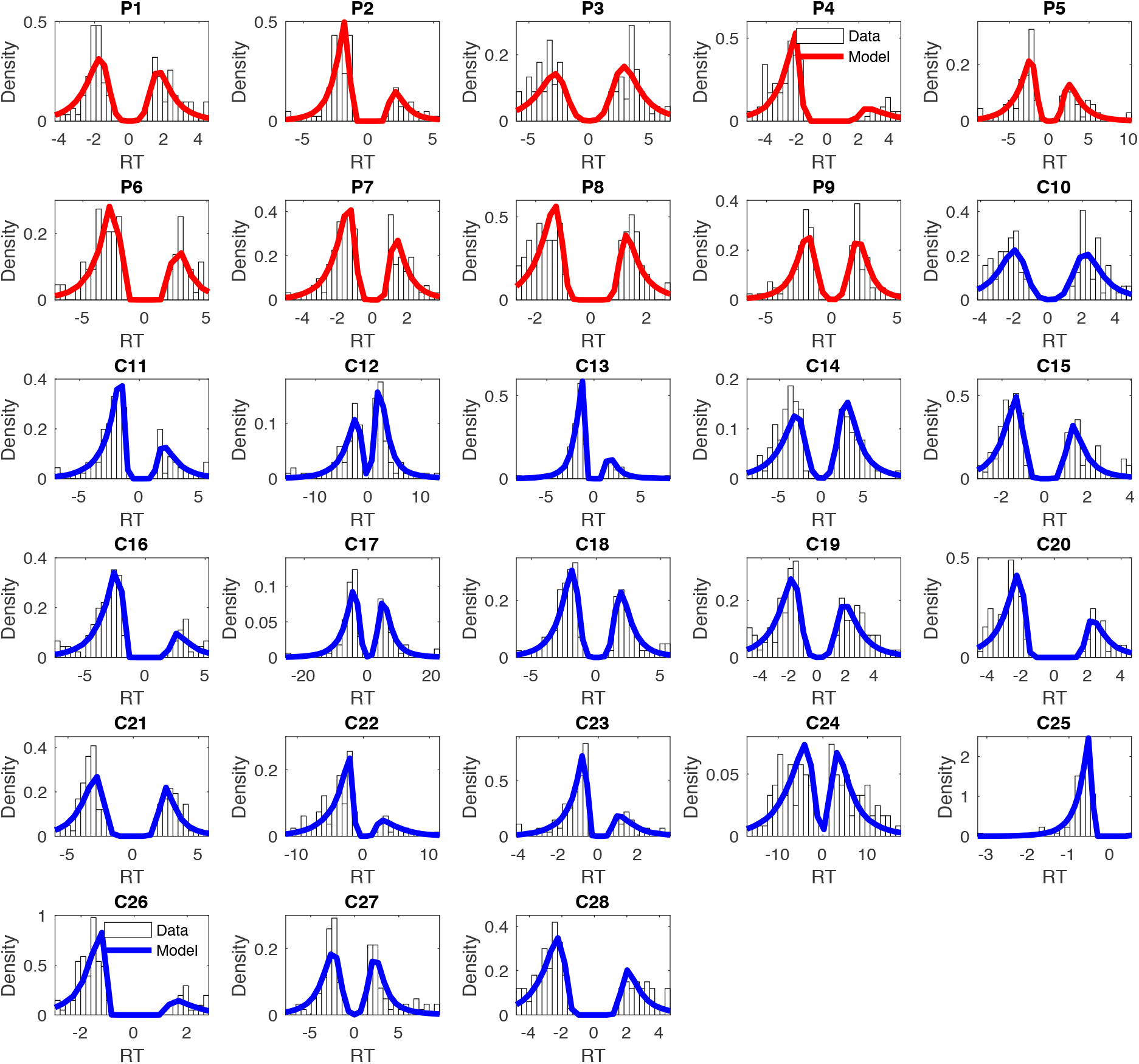
Posterior predictive plots of the drift diffusion probability discounting / risky choice model with non-linear value scaling of the drift rate (DDM_S_) for all participants (red – mOFC patients, blue – controls). Histograms depict the observed RT distributions for each participant. The solid lines are smoothed histograms of the model predicted RT distributions from 1000 individual subject data sets simulated from the posterior of the winning hierarchical model. RTs for choices of the safe option are plotted as negative, whereas RTs for risky choices are plotted as positive. The x-axes are adjusted to cover the range of observed RTs for each participant.

We next checked the degree to which the best-fitting models predicted participants’ binary choices. To test this, we used each participant’s mean posterior parameters from the hierarchical model to calculate model predicted choices, and compared these to the observed choices. Median accuracy was >.90 for all tasks and groups (see Table 4), suggesting that the best-fitting hierarchical models captured individual choices similarly well in both groups and tasks.

**Table 4.** Median (range) of the proportion of correctly predicted binary choices of the DDM_S_ for each task and group.

|  | DDM<sub>S</sub>: Temporal discounting | DDM<sub>S</sub>: Risk taking |
| --- | --- | --- |
| mOFC patients | .92 (.89-.99) | .92 (.84-.98) |
| Controls | .91 (.84-.99) | .91 (.82-.99) |

### Model validation

With the drift diffusion model as the choice rule, we introduced additional complexity, as softmax action selection typically only has a single free parameter (*β*). Therefore, we next examined the correspondence of choice model parameters (i.e. *log(k)*_*now*_, *shift*_*log(k)*_ and *log(h)*) between models estimated using softmax vs. DDM_S_ choice rules. One would not expect preferences as reflected in these parameters to differ systematically as a function of whether only binary choices are fitted (softmax) or whether both choices and reaction times are jointly fitted (DDM_S_). A convergence between the estimated parameters from the two choice rules would therefore strengthen confidence in the applicability of the DDM in the context of the present tasks.

To this end, we extracted the mean single-subject parameter estimates for *log(k)*_*now*_ (the hyperbolic discount rate in the *now* condition of the temporal discounting task, Eq. 1), *shift*_*log(k)*_ (the parameter modeling the reduction in discounting between *now* and *not now* conditions in the temporal discounting task, Eq. 1) and *log(h)* (reflecting the degree of discounting of value over probabilities, Eq. 2) from the hierarchical fits of the two winning DDM_S_ models as well as from the hierarchical fits using standard softmax action selection. Figure 4 shows scatter plots of mean single-subject parameters estimated via softmax vs. via DDM_S_. Correlations were very high between the different choice rules (temporal discounting: *log(k)*_*now*_ *r*=.93, *shift*_*log(k)*_ *r*=.91; risky choice/probability discounting: *log(h) r*=.98). Since the correlation for *shift*_*log(k)*_ appeared to be somewhat inflated by the extreme datapoints of the mOFC patients, we re-ran the correlation only in the control group. Here, the correlation was lower but still robust (*r*=.52). Together, these analyses confirm that parameters estimated via softmax modeling of binary choices can be reliably reproduced when jointly fitting choices and RTs via the DDM_S_.

**Figure 4.**
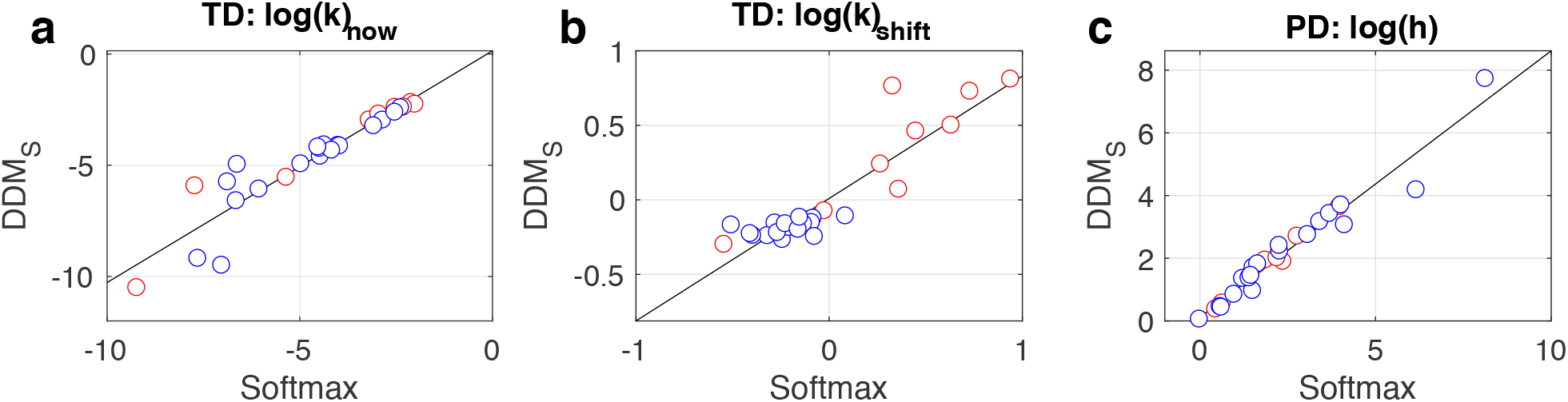
Consistency of model parameters for temporal discounting (TD: a/b) and probability discounting (PD, c) between softmax and DDM_S_ choice rules. Scatter plots (controls: blue, mOFC patients: red) show model parameters estimated via a standard softmax choice rule (x-axis) vs. parameters estimated via a drift diffusion model choice rule with non-linear drift rate scaling (DDM_S_, y-axis). a) Temporal discounting log(discount rate) for *now* trials. b) Shift in log(k) between *now* and *not now* trials). c) Probability discounting log(discount rate).

Along similar lines, we next checked estimated DDM parameter against model-free RT statistics (minimum and median RT). The non-decision time *τ* captures RT components unrelated to the evidence accumulation process, and therefore reflects individual differences related to e.g. perceptual processing of the decision options and/or response preparation and execution. That is, for *τ* one would predict positive correlations in particular with the minimum RT, and to a lesser extent with median RT. As expected, correlations of *τ* with minimum and median RT where significant and more pronounced for minimum RT (Figure 5a,c: temporal discounting *r*_minRT_=.95, *r*_medianRT_=.69; Figure 5b,d: probability discounting *r*_minRT_=.92, *r*_medianRT_=.54).

**Figure 5.**
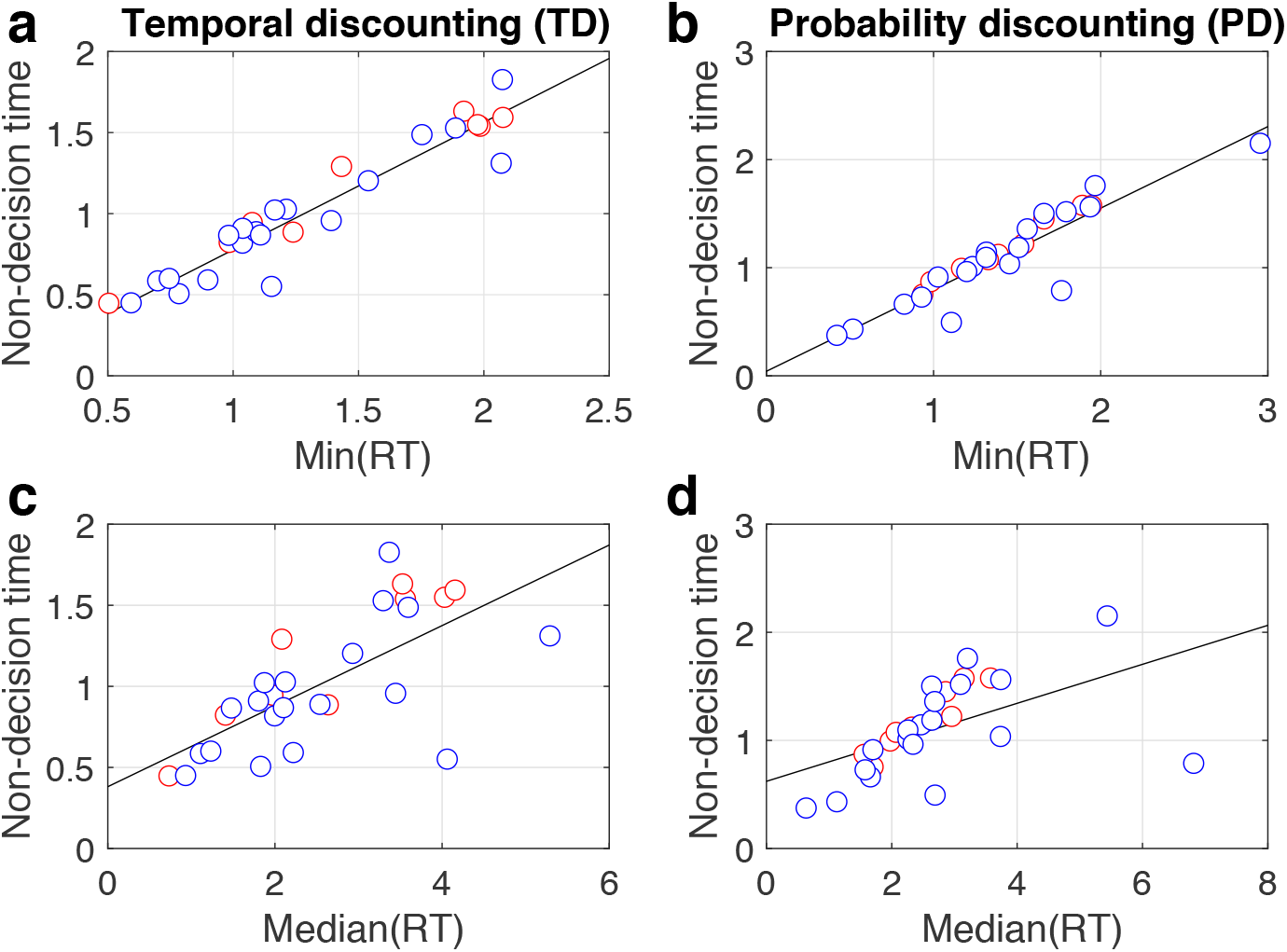
Scatter plots (controls: blue, mOFC patients: red) showing correlations between model-based non-decision time from the best fitting DDM_S_ models (x-axis) and minimum RT (a/b) and median RT (c/d) for temporal discounting (a/c) and risky choice / proability discounting (b/d).

The boundary separation parameter *α* on the other hand reflects the threshold that the accumulated evidence needs to exceed before participants commit to a decision. Again one would expect positive correlations with minimum and median RT, but a more pronounced association with median RT. This is exactly what we observed (Figure 6a,c: temporal discounting *r*_minRT_=.71, *r*_medianRT_=.94; Figure 6b,d: probability discounting *r*_minRT_=.67, *r*_medianRT_=.88).

**Figure 6.**
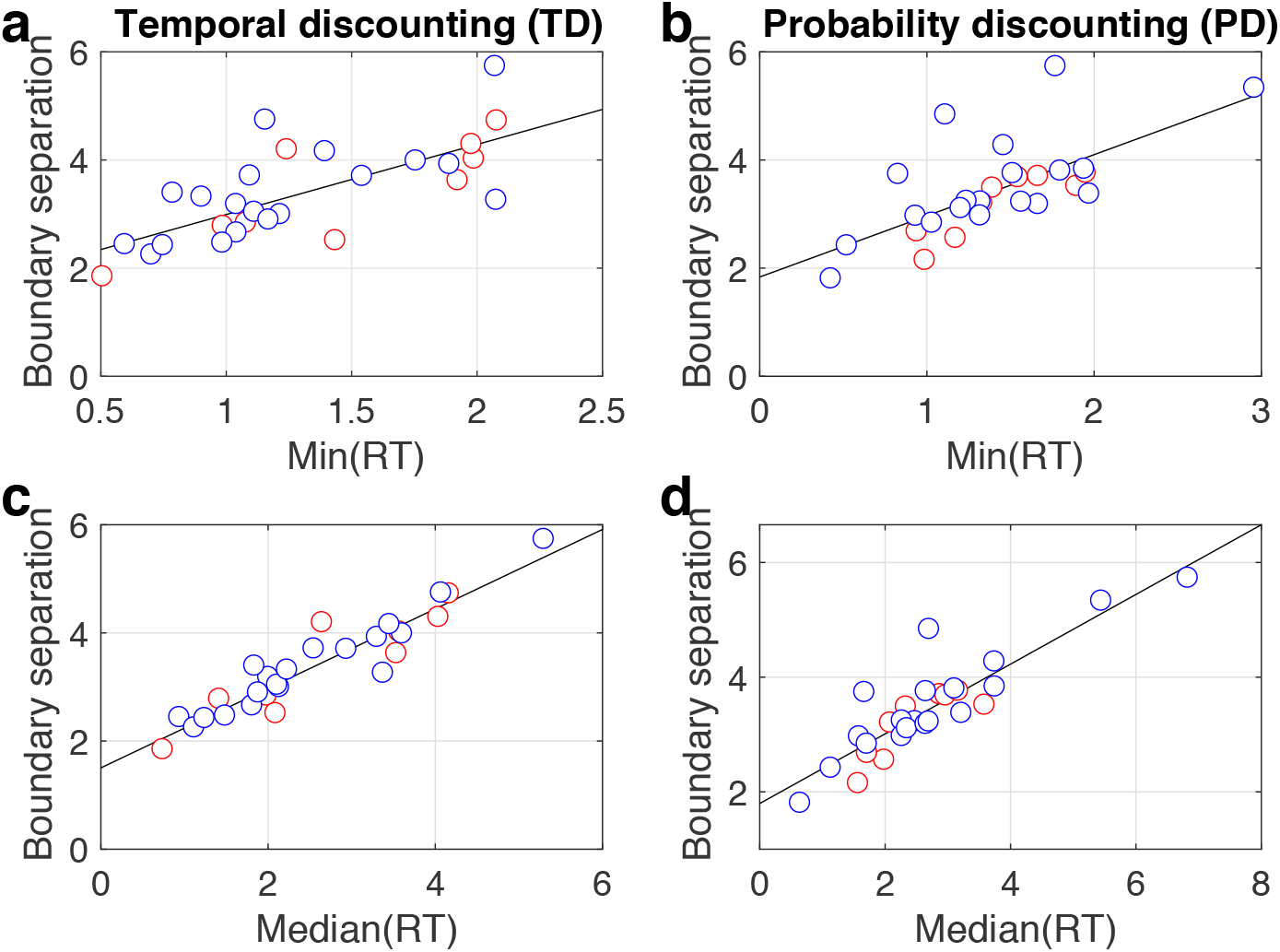
Scatter plots (controls: blue, mOFC patients: red) showing correlations between model-based boundary separation parameter from the best fitting DDM_S_ models (x-axis) and minimum RT (a/b) and median RT (c/d) for temporal discounting (a/c) and risky choice / proability discounting (b/d).

### Comparison to previous model-free analyses in mOFC patients

We have previously reported that temporal discounting in mOFC lesion patients is more affected by the immediacy of SS rewards than in controls^31^. Our previous analysis revealed this both via an analysis of the area-under-the-curve of the empirical discounting function^57^ and by a direct comparison of preference reversals between groups. To further validate the applicability of the DDM in the context of temporal discounting, we next examined whether these effects could be reproduced via the hierarchical DDM_S_. Figure 7 plots the group-level posterior distributions of parameter means for all seven parameters, where we for the purposes of comparison to our previous results first focus on *log(k)*_*now*_ (the discount rate in the baseline *now* condition, see Figure 7f) and *shift*_*log(k)*_ (the parameter modeling the *decrease* in discounting in *not now* trials as compared to *now* trials, see Figure 7g). The analysis of directional between-subject effects revealed a numerical increase in *log(k)*_*now*_ in the mOFC patient group (Figure 7f, Table 5) and strong evidence for a substantially greater difference in discounting between *now* and *not now* trials in the mOFC patient group (Figure 7g, Table 3). This shows that our results based on model-free summary measures of discounting behavior following mOFC lesions^31^ could be reproduced via a hierarchical Bayesian estimation scheme with the DDM_S_ as the choice rule.

**Table 5.**
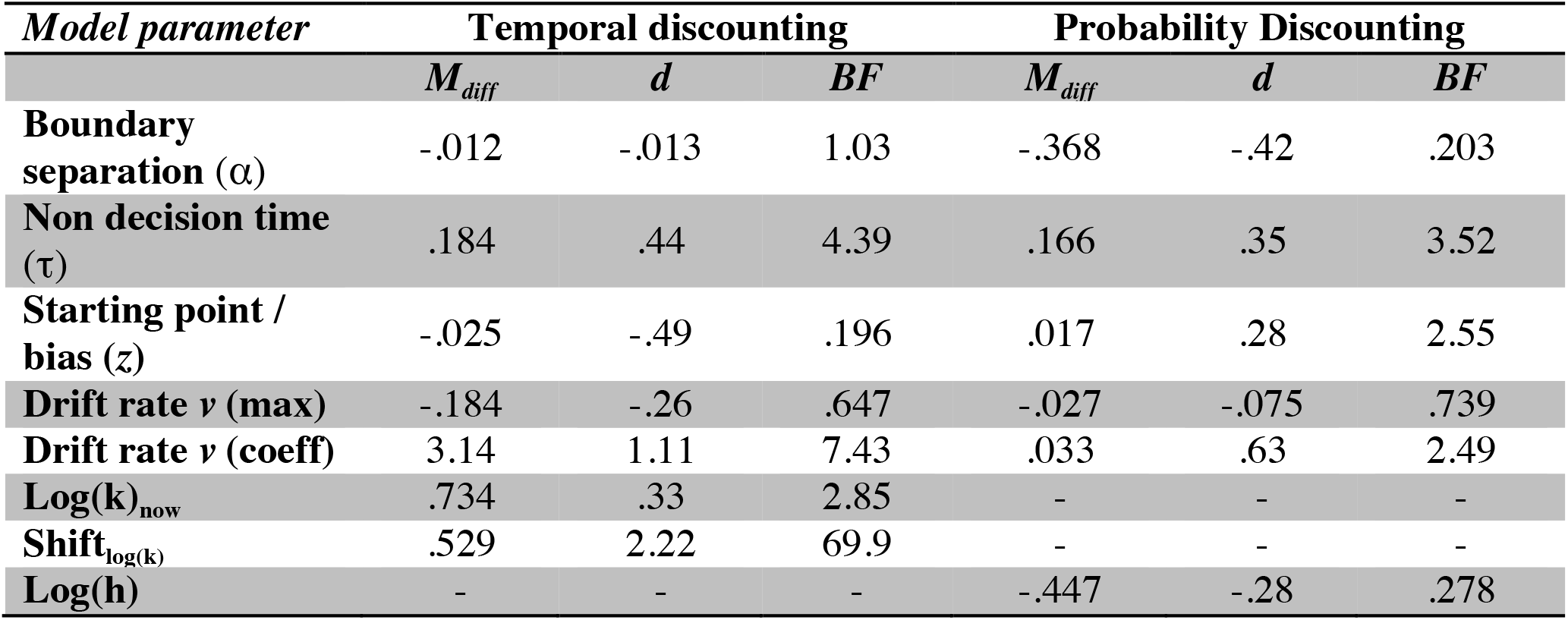
Summary of group differences in model parameters. For each parameter and task, we report the mean difference in the group-level posterios (M_diff_: patients – controls) and Bayes Factors testing for directional effects^14,55^. Bayes Factors <.33 indicate evidence for a reduction in the patient group, whereas Bayes Factors >3 indicate evidence for an increase in the patient group (see Methods section). Standardized effect sizes (Cohen’s *d*) were calculated based on the posterior group-level estimates of mean and precision (see methods section).

| *Model parameter* | **Temporal discounting** |  |  | **Probability Discounting** |  |  |
| --- | --- | --- | --- | --- | --- | --- |
| | $M_{\text{diff}}$ | $d$ | $BF$ | $M_{\text{diff}}$ | $d$ | $BF$ |
| **Boundary separation ($\alpha$)** | -.012 | -.013 | 1.03 | -.368 | -.42 | .203 |
| **Non decision time ($\tau$)** | .184 | .44 | 4.39 | .166 | .35 | 3.52 |
| **Starting point / bias ($z$)** | -.025 | -.49 | .196 | .017 | .28 | 2.55 |
| **Drift rate $v$ (max)** | -.184 | -.26 | .647 | -.027 | -.075 | .739 |
| **Drift rate $v$ (coeff)** | 3.14 | 1.11 | 7.43 | .033 | .63 | 2.49 |
| **Log($k$)<sub>now</sub>** | .734 | .33 | 2.85 | - | - | - |
| **Shift<sub>log(k)</sub>** | .529 | 2.22 | 69.9 | - | - | - |
| **Log($h$)** | - | - | - | -.447 | -.28 | .278 |

**Figure 7.**
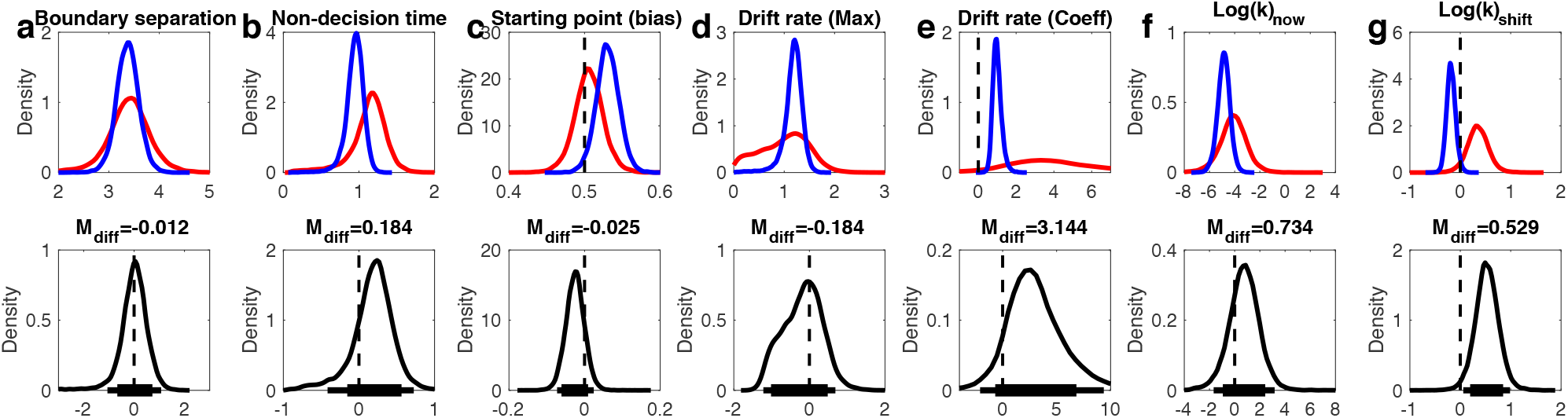
Modeling results for the DDM_S_ temporal discounting model. Top row: posterior distributions of the parameter group means for controls (blue) and mOFC patients (red). Bottom row: Posterior group differences (mOFC patients – controls) for each parameter. Solid horizontal lines indicate highest density intervals (HDI, thick lines: 85% HDI, thin lines: 95% HDI).

### *Risk-taking in* vmPFC/mOFC *patients*

Risk-taking on the probability discounting task was quantified via the probability discounting parameter *log(h)*, where higher values indicate a greater discounting of value over probabilities. There was some evidence for a smaller *log(h)* in vmPFC/mOFC (Figure 8f, Table 5), reflecting a relative increase in risk-taking (reduced value discounting over probabilities) as compared to controls.

**Figure 8.**
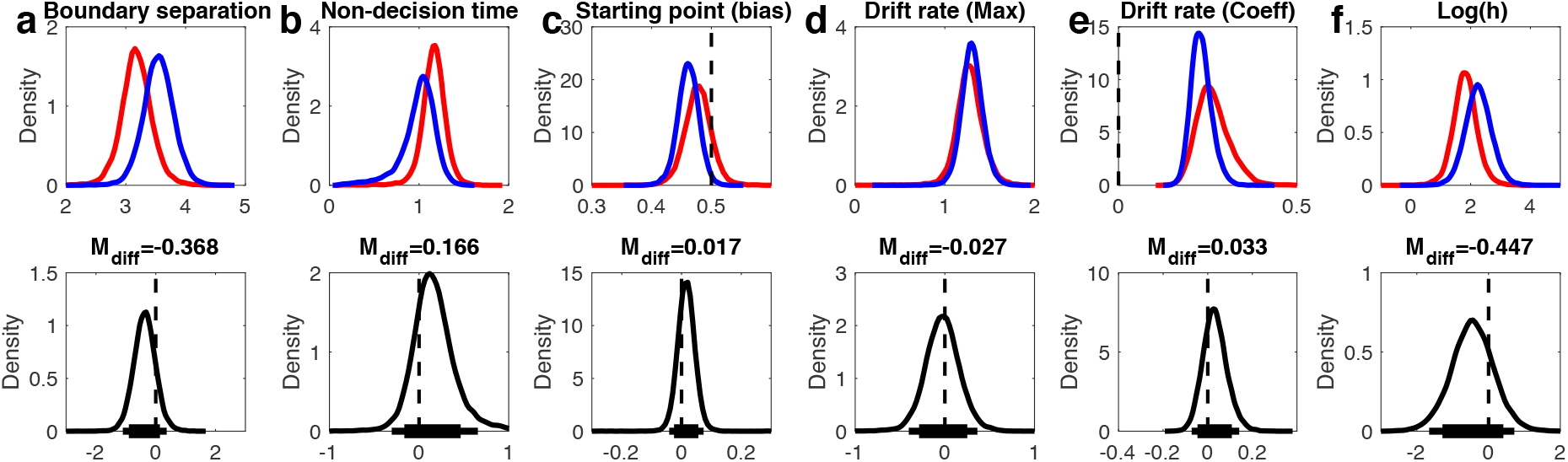
Modeling results for the DDM_S_ risky choice model. See Figure 7 for details.

### Effects of mOFC lesions on diffusion model parameters

Finally, we examined the diffusion model parameters of the DDMS models in greater detail. First, there was evidence for substantially longer non-decision times in the vmPFC/mOFC patient group for both tasks (see Table 5 and Figures 7b, 8b). These effects amounted to on average 184ms for temporal discounting and 166ms for risky choice. Second, the group differences observed for the starting point (bias) parameter largely mirrored group differences observed for discounting behavior. For temporal discounting, controls exhibited a more pronounced bias towards the LL boundary than vmPFC/mOFC patients, who exhibited a largely neutral bias here. For risky choice, controls showed a bias that was numerically shifted towards the safe-option compared to vmPFC/mOFC patients. Third, posterior distributions for the boundary separation parameter (alpha) in temporal discounting showed high overlap and the difference distribution was centered at 0 (Figure 7a). In contrast, for risky choice, there was evidence for a reduced boundary separation in the vmPFC/mOFC patients (Figure 8a, Table 3).

In the DDM_S_, two components of the drift rate can be dissociated: the asymptote of the drift rate scaling function (*v*_*max*_), that is, the maximum drift rate that is approached as value differences increase, and the slope of the sigmoid (*v*_*coeff*_). In both tasks, there was no evidence for a group difference in *v*_*max*_ (see Table 5 and Figures 7d, 8d) and both difference distributions were centered at 0. Across tasks and groups, the value scaling parameter for the drift rate (*v*_*coeff*_) was generally > 0, reflecting a robust positive effect of value differences on the rate of evidence accumulation (see Figure 7d, 8d). Interestingly, the drift rate slope parameter (*v*_*coeff*_) was increased in the vmPFC/mOFC patients for both tasks, an effect that was substantial for the temporal discounting data. Here, the posterior distribution also had a much higher variance compared to the control group. This was driven by 4/9 vmPFC/mOFC patients who had *v*_*coeff*_ estimates that fell substantially beyond the values observed in controls and in the remaining patients (mean *v*_*coeff*_ estimates: P1: 17.89, P3: 8.32, P4: 3.38, P5: 4.70). These extreme cases included the patient with the lowest discount rate (P1 *log(k)*_*now*_: −10.53) and the patient with the second highest discount rate (P4 *log(k)*_*now*_: −2.28).

## Discussion

We compared different choice rules for modeling inter-temporal and risky choice / probability discounting in healthy controls and patients with vmPFC/mOFC lesions. For each task we examined a standard softmax action selection function and three variants of the drift diffusion model (DDM). Across tasks, the data were better accounted for by a DDM with a non-linear mapping of value differences onto the drift rate (DDM_S_) than by a DDM with linear mapping (DDM_lin_) or a null model without any value modulation (DDM_0_). In a series of further model checks, we verified 1) the consistency between choice model parameters estimated via the DDM_S_ as compared to a standard softmax choice rule, 2) the correspondence between observed RT distributions and RT distributions simulated from the estimated posterior distributions (posterior predictive checks) for each individual participant and 3) the expected associations between the DDM parameters non-decision time and boundary separation and non-parametric RT measures. We then used the DDM_s_ to reproduce our previous results on temporal discounting in patients with vmPFC/mOFC lesions^31^, to characterize risk-taking behavior in these patients, and to explore group differences in diffusion model parameters across tasks.

Previous studies have successfully incorporated RTs in the modeling of value-based decision-making, e.g. via the linear ballistic accumulator model^16^ or linear regression^13^. Here we build on recent work in reinforcement learning^12,14,15^ and examined the degree to which the drift diffusion model could be used as the choice rule in temporal discounting and risky choice. In line with a recent model comparison in reinforcement learning^15^, our model comparison of linear vs. non-linear value scaling revealed a superior fit of the DDM with non-linear (sigmoid) value scaling both for temporal discounting and risky choice data. Posterior predictive checks of the winning models revealed a reasonably good fit to the observed RT distributions of most individual participants. However, in a few participants, models tended to underestimate RTs for choices towards the lower boundary, and this was perhaps most evident in the case of the risky choice data in some of the vmPFC/mOFC patients.

One advantage of hierarchical Bayesian parameter estimation is that robust model fits can be obtained with fewer data points than are typically required for maximum likelihood estimation^58,59^, and this is also the case for the drift diffusion model^58^. The reason is that in contrast to obtaining single-subject point estimates of parameters (as in maximum likelihood estimation), in hierarchical Bayesian estimation, the group-level distribution of parameters constrains and informs the parameters estimated for each participant. One consequence of this is shrinkage^59^ or partial pooling, such that in a hierarchical model individual parameter estimates tend to be drawn towards the group mean. While this can improve the predictive accuracy of parameters, there is the possibility that meaningful between-subjects variability is removed^60^. Nonetheless, we believe that for situations with limited data per subject^58^, which is a particular issue in studies involving lesion patients, the hierarchical Bayesian estimation approach is most appropriate.

We examined variants of the DDM in tasks where they have not been applied previously (although other sequential sampling models have^16^). We therefore ran a number of additional analyses to validate our modeling results. First, we checked whether discount rates estimated via softmax and via the DDM_S_ showed robust correlations. They did. Although this might at first glance seem like an obvious result, it is nonetheless reassuring that parameter estimates obtained via standard methods can be reliably reproduced using the DDM_S_. Along similar lines, we checked the correlation of non-decision time and boundary separation parameters from the DDM_S_ with model-free RT summary statistics. Again, we observed the expected associations (Figure 6,7), suggesting that the results obtained via the DDM_S_ are valid. Finally, our analysis of the DDM_S_ for temporal discounting reproduced our previous model-free results in vmPFC/mOFC patients^31^: discounting behavior following vmPFC/mOFC damage was substantially more affected by *SS* reward immediacy than in controls, which in the present modeling scheme was reflected in a substantially increased *shift*_*log(k)*_ parameter in the vmPFC/mOFC patient group. Together, these observations strengthen our confidence in the validity of using the DDM as the choice rule in inter-temporal and risky choice.

The stimulus coding scheme that we adopted here differs from accuracy coding as implemented in recent applications of the DDM to reinforcement learning^14,15^, with implications for the interpretation of the DDM parameters. First, the drift rate *v* in the present coding scheme (as reflected in *v*_*max*_ and *v*_*coeff*_) can be interpreted along similar lines as in classical perceptual decision-making tasks: it reflects the rate of evidence accumulation. In stimulus coding, however, higher drift rates do not directly correspond to better performance (as is the case in accuracy coding), because there is no objectively correct response. Instead the drift rate parameters reflect a participant’s overall sensitivity to value differences, similar to inverse temperature parameters in softmax models. Second, in accuracy coding, the boundary separation *α* governs the speed-accuracy trade-off^9^, such that a larger boundary separation corresponds to a focus on accuracy rather than speed. In the absence of “accuracy”, boundary separation in stimulus coding reflects similarly the amount of evidence required to commit to a decision, but here response caution^10^ might be a more appropriate term. Finally, adopting stimulus coding allowed us to estimate a starting point (bias) parameter. In all cases, the estimated bias parameters were relatively close to 0.5 (a neutral bias). Nonetheless, the group differences in bias that we observed for each task mirrored the results for the choice model parameters. That is, the group that displayed a preference for one option as reflected in the discount rate parameter (e.g. LL rewards in the case of controls) also exhibited a response bias towards that decision boundary.

Our results also provide novel insights into the role of the vmPFC/mOFC in decision-making. Using a model-based analysis, we show that value-differences exert a similar (if not stronger) effect on trial-wise drift rates in vmPFC/mOFC patients compared to controls, whereas the maximum drift rate *v_max_* was of similar magnitude in the two groups. This is in line with an earlier report showing reduced preference consistency but no changes in overall RTs or the value-modulation of RTs in vmPFC/mOFC patients^41^. If one considers the overwhelming evidence of neuroimaging studies showing a prominent role of the vmPFC/mOFC in reward valuation^43,44^, it is nonetheless striking that lesions to this region do not negatively impact the value-sensitivity of the evidence accumulation process during value-based decision-making. Our data are therefore more compatible with the idea that vmPFC/mOFC, likely in interaction with other regions^61,62^, plays a role in self-control, such that lesions shift preferences towards options with a greater short-term appeal. Previous work has suggested that damage to vmPFC/mOFC might decrease the temporal stability of value representations, leading to inconsistent preferences^40–42^. There was no evidence in the present data that the lesion patients’ decisions were more “noisy” or “erratic”. Similar to a previous study on temporal discounting^32^, choice consistency was high such that the best-fitting DDMS accounted for about 90% of choices in both groups and tasks. Together with the intact value modulation of the drift rate, this suggests that value representations on a given trial^41^ and throughout the course of testing sessions were relatively stable in both groups. In contrast, results from both tasks revealed a substantial increase in non-decision times in the patient group. Together, these observations suggest that vmPFC/mOFC lesions lead to a slowing of more basic perceptual and/or response-related processes during value-based decision-making, while leaving the effects of value-differences on the evidence accumulation process strikingly intact.

Previous studies have shown increases in risky decision-making following vmPFC/mOFC damage^34,36^. Our finding of attenuated discounting over probabilities in vmPFC/mOFC patients is consistent with these previous results. However, our model-based analysis revealed an additional effect: lesion patients also exhibited reduced response caution during risky choice, reflected in a reduced boundary separation parameter. In contrast, this was not observed for temporal discounting. This suggests that risk taking in vmPFC/mOFC patients might not only be driven by altered preferences, but also by more premature responding.

Taken together, our results demonstrate the feasibility of using the drift diffusion model as the choice rule in the context of inter-temporal and risky decision-making. Model comparison revealed that a variant of the DDM that included a non-linear drift rate modulation provided the best fit to the data. We further show that choice model parameters estimated via the DDM show a close correspondence to parameters estimated via standard methods. Finally, the application of a sequential sampling model revealed additional insights: while the value-dependency of the evidence accumulation process was strikingly unaffected by vmPFC/mOFC damage, we observed a slowing of non-decision times both in temporal discounting and risky choice, with implications for models of decision-making. This modeling framework might provide further insights, e.g. when studying mechanisms underlying context-dependent changes in decision-making^63–66^ or impairments in decision-making in psychiatric^67^ and neurological disorders^6^.

## Acknowledgements

J.P. and M.D. conceived research. J.P. performed research, analyzed the data and wrote the paper. M.D. provided critical revisions. This work was funded by Deutsche Forschungsgemeinschaft (grants PE1627/4-1 and PE1627/5-1 to J.P.). We thank Donatella Scabini for help with patient recruitment, Natasha Young for help with testing control subjects and members of the Peters Lab for helpful discussions.

